# Restoration of deficient DNA Repair Genes Mitigates Genome Instability and Increases Productivity of Chinese Hamster Ovary Cells

**DOI:** 10.1101/2021.01.07.425558

**Authors:** Philipp N. Spahn, Xiaolin Zhang, Qing Hu, Nathaniel K. Hamaker, Hooman Hefzi, Shangzhong Li, Chih-Chung Kuo, Yingxiang Huang, Jamie C. Lee, Peter Ly, Kelvin H. Lee, Nathan E. Lewis

**Affiliations:** Department of Pediatrics, University of California, San Diego, La Jolla, CA 92093; The Novo Nordisk Foundation Center for Biosustainability at the University of California, San Diego School of Medicine, San Diego, La Jolla, CA 92093; Department of Chemical and Biomolecular Engineering, University of Delaware, Newark, DE 19711; Department of Pathology, University of Texas Southwestern Medical Center, Dallas, TX 75390; Department of Bioengineering, University of California, San Diego, La Jolla, CA 92093

**Keywords:** Genome instability, systems biology, cell line instability, CHO, DNA repair, biotechnology

## Abstract

Chinese Hamster Ovary (CHO) cells are the primary host used for manufacturing of therapeutic proteins. However, production instability of high-titer cell lines is a major problem and is associated with genome instability, as chromosomal aberrations reduce transgene copy number and decrease protein titer. We analyzed whole-genome sequencing data from 11 CHO cell lines and found deleterious single-nucleotide polymorphisms (SNPs) in DNA repair genes. Comparison with other mammalian cells confirmed DNA repair is compromised in CHO. Restoration of key DNA repair genes by SNP reversal or expression of intact cDNAs improved DNA repair and genome stability. Moreover, the restoration of LIG4 and XRCC6 in a CHO cell line expressing secreted alkaline phosphatase mitigated transgene copy loss and improved protein titer retention. These results show for the first time that correction of key DNA repair genes yields considerable improvements in stability and protein expression in CHO, and provide new opportunities for cell line development and a more efficient and sustainable production of therapeutic proteins.

## Introduction

Chinese Hamster Ovary (CHO) cells have been the leading expression system for the industrial production of therapeutic proteins for over 40 years, and projections show they will maintain this dominant position into the foreseeable future because they produce >80% of therapeutic proteins approved between 2014-2018 corresponding to more than $100 billion of products per year [1, 2]. Steady improvements in cell line development, media formulation, and bioprocessing now enable production yields exceeding 10 g/L from a fed-batch culture. Emerging resources, including the CHO and hamster genome sequence [3–8] and genome editing tools [9–14] now allow researchers to rely less on largely empirical, “trial-and-error” approaches to CHO cell line development, and move towards a more rational engineering approach, in pursuit of novel CHO lines with tailored, superior attributes [15–18].

However, cell line instability, i.e. the propensity of a cell line to lose industrially relevant properties over time, remains a challenging problem in the field as it can result in loss of titer, changes in product quality, or changes in cell growth potentially impacting manufacturing and drug supply. Thus, considerable effort is expended to ensure that production clones are stable for use in fed-batch production (for example, that a given clone retains 70% of its titer after 60 generations in culture which would be deemed a “stable” producer [19]). However, as the field moves toward perfusion and continuous culture, cell line stability remains a key challenge to overcome.

Production instability can arise from transcriptional transgene silencing through epigenetic mechanisms, such as promoter methylation or histone acetylation [20–26]. However, while such epigenetic transcriptional silencing was associated with production instability in some studies, a loss in transgene copy number due to genome instability (i.e. the rapid accumulation of mutations and chromosome aberrations) appears to be predominant cause of production instability [20, 22, 27–33]. The inability to maintain genome integrity will negatively affect production stability in most, if not all, production cell lines during long-term culture. Because massive transgene expression imposes a high metabolic demand on the host, the random loss of these transgenes is effectively under positive selection in the bioreactor, causing the emergence of non-producing subpopulations that can outgrow the remaining producers in the pool, resulting in titer decline.

Genome instability is common across cancers, and can arise from defects in DNA damage repair [34]. In CHO, genome instability was first reported in the 1970s when direct observations of CHO chromosomes revealed a divergence from the Chinese hamster (*Cricetulus griseus*) karyotype and variability in karyotypes among CHO clones [35, 36]. These karyotype variations occur regardless of growth condition, and do not differ markedly between pooled and clonal populations [37–39]. Loss of chromosomal material and improper chromosome fusions (translocations) are caused by improperly repaired double-strand breaks (DSBs) [40, 41] which result from metabolic processes, such as attack by free radicals or collapsed DNA replication forks [41]. Given that primary cells from the Chinese hamster do not show genome instability, DSB repair ability is likely compromised in CHO. Thus, identifying and restoring deficient DSB repair genes in CHO cells could effectively enhance the DSB repair system, mitigate genome instability, and improve product titer stability.

To date, few studies have addressed the persistent genome instability problem in CHO cells, and it has been challenging to develop effective counterstrategies. Detailed quantification of chromosomal instabilities in production cell lines has indicated that certain chromosome sites are less prone to instability than others [42]. This observation has suggested that transgene loss may be avoided by targeting transgenes to these stable chromosomal areas, an option now possible through the development of targeted transgene integration techniques [12, 43–45]. Further studies used gene knock-outs (*ATR* and *BRCA1*, respectively) to increase product titer by increasing transgene copy number amplification [46, 47], but whether these knock-outs are able to sustain high production in long-term culture has remained unclear. Thus, a pressing need remains for novel approaches to mitigate production instability stemming from DSBs. Although the mechanistic connections between production instability and DNA DSBs are evident, the field has not systematically explored the engineering of DNA repair as a possible means to reduce transgene loss in CHO. Inactivation of *ATR* was shown to result in an increase in transgene copies during the amplification phase, but also a less rigid cell cycle control and higher chromosomal instability [46], which might exacerbate production instability in the long run. Therefore, rather than inactivating DNA repair genes for short-term gains, rescue of deficient DNA repair could constitute a promising approach to achieve long-term improvement in stability.

Here we investigate the relationship between DSB repair and genome instability in CHO cells. We describe the landscape of mutations in DNA repair genes across a panel of 11 commonly used CHO lines, and show that DSB repair ability is compromised in CHO compared to other mammalian cell lines. We then show that restoration of key DSB repair genes yields significant improvements in repair ability and genome stability. Finally, we show in a proof-of-concept experiment that restoration of DNA repair genes *XRCC6* and *LIG4* improves the stability of transgene copy number and protein titer in a long-term-culturing study in a CHO cell line secreting alkaline phosphatase (SEAP).

## Results

### CHO cells are deficient in DSB repair

To test if DSB repair is deficient in CHO cells, we compared DSB repair between CHO-K1 and bEnd.3 and BHK-21, two established and well characterized mammalian cell lines. An endogenous DSB level was first estimated by counting the number of γH2AX foci per cell. γH2AX denotes phosphorylated histone H2AX in the chromatin area surrounding a DSB which often extends several megabases from the break site, visible as a focus in confocal microscopy [48]. Quantification of γH2AX foci is a well-established method to visualize unrepaired DSBs as H2AX is dephosphorylated only after initiation of repair [49]. The number of endogenous DSBs in CHO-K1 was significantly higher than in BHK-21 cells (Fig. 1b,c: untreated). CHO cells also had more endogenous DSBs than bEnd.3 cells, although the difference was less pronounced (Fig. 1b,c: untreated). We next tested the repair ability in these cell lines after treatment with bleomycin, a DSB-inducing drug. A 12-hour treatment (10 μg/mL) allowed sufficient passive diffusion across the cellular membrane, and remaining DSBs were counted at different time-points during the recovery period (Fig. 1a). bEnd.3 cells showed the fewest DSBs with an average of 12 foci per nucleus 1h after bleomycin removal, and a reduction to an average of 1.5 foci per nucleus after 24h (Fig. 1b,c; 1.0 DSBs/hour by linear regression). BHK-21 cells showed comparable DNA damage to CHO cells 1h after bleomycin removal, but a fast reduction to an average of 4 foci per nucleus after 24h (Fig. 1b,c; 2.6 DSBs/hour by linear regression). CHO-K1 showed the slowest DSB repair rate (Fig. 2b,c; 0.9 DSBs/hour by linear regression) and the most remaining DSBs after 24h of recovery, showing that DSB repair is compromised in CHO cells.

**Fig 1:**
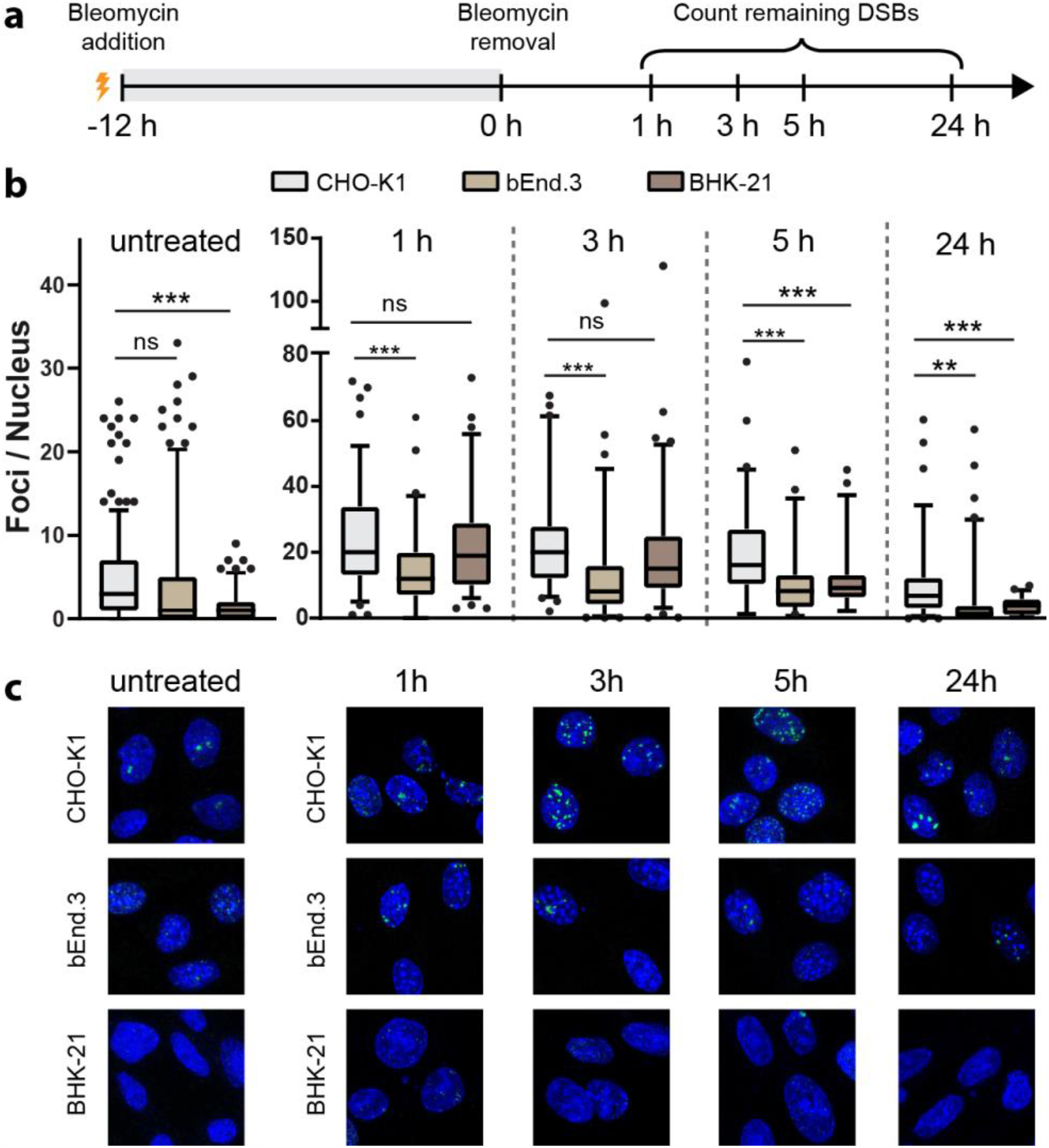
DSB repair ability in CHO cells and primary cells. **(a):** Method of bleomycin treatment: Cells were incubated in 10 μg/mL bleomycin for 12 hours, after which bleomycin was removed, wells were rinsed with DPBS, and cells were left in full growth medium without bleomycin for the indicated time periods for recovery before fixation. **(b):** Enumeration of γH2AX foci in CHO-K1, bEnd.3, and BHK-21 cells without bleomycin treatment (left) and after bleomycin treatment and the indicated recovery intervals. Welch’s t-tests. untreated: n≥129 nuclei, 1h: n≥63 nuclei, 3h: n≥66 nuclei, 5h: n≥45 nuclei, 24h: n≥ 42 nuclei. Whiskers showing 5/95-percentiles. **(c)**: Immunostainings against γH2AX in CHO-K1 wildtype, bEnd.3, and BHK-21 cells, corresponding with the conditions in (b). Cells counterstained with DAPI.

**Fig 2:**
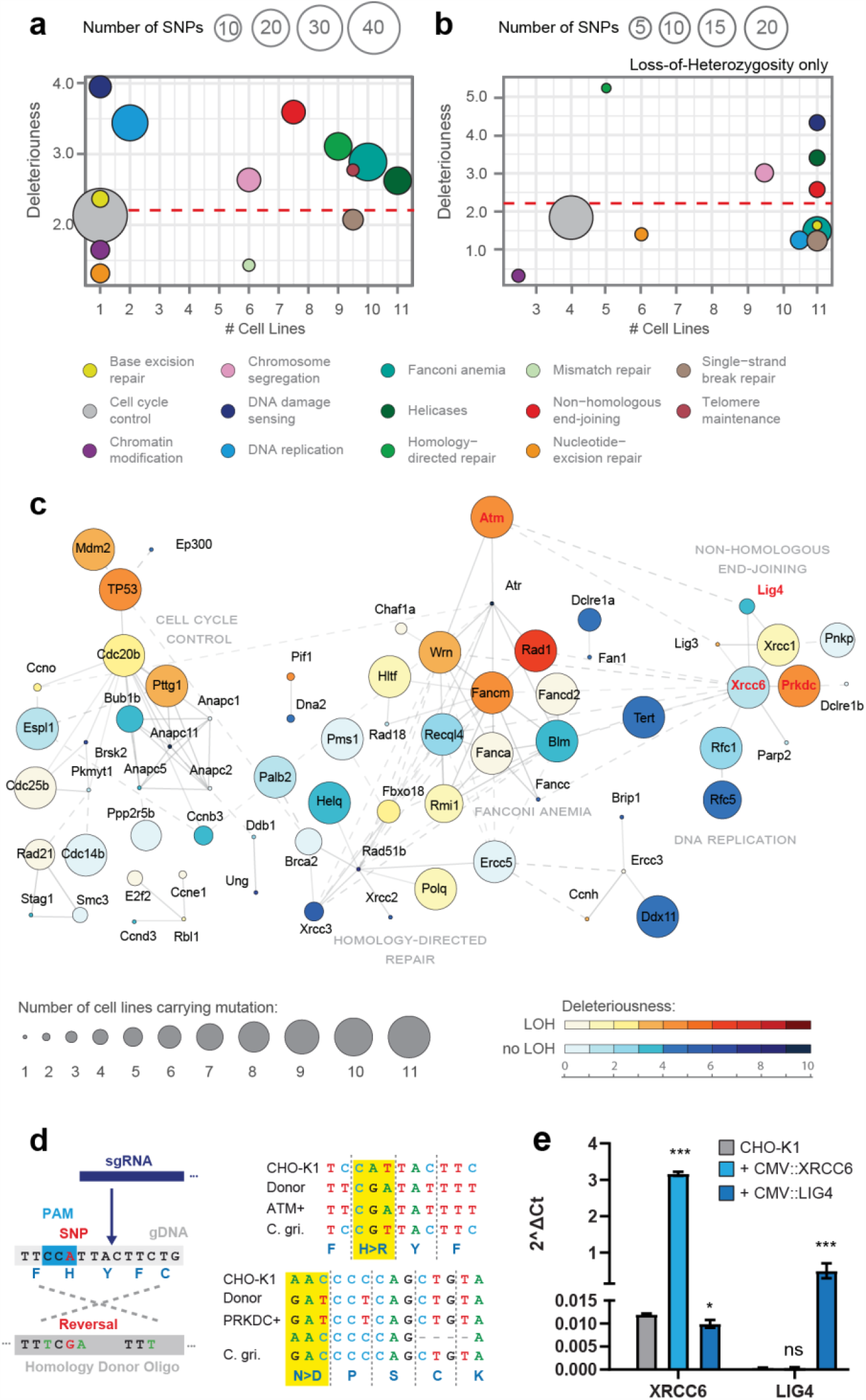
DNA repair mutation landscape in CHO. **(a):** Mutations found in whole-genome sequencing data from 11 major CHO cell lines, categorized into 14 gene ontology groups (bottom legend) related to genome stability. The number of CHO lines carrying each mutation and the predicted deleteriousness are averaged across all mutations in each ontology group (x-axis: median of cell lines affected; y-axis: average negative PROVEAN score). The dashed line indicates the recommended threshold (2.282) to separate neutral from detrimental SNPs [51]. In total, 157 mutations are shown. **(b):** Subset of mutations shown in (a) that have undergone loss of heterozygosity, i.e. having lost the Chinese hamster wildtype allele at the respective locus (62 mutations total). **(c)**: Protein-protein interaction network of genes carrying mutations in at least one CHO cell line. Number of CHO cell lines carrying each mutation is displayed by bubble size. Color represents predicted deleteriousness (negative PROVEAN score) and loss of heterozygosity (LOH). In cases where multiple mutations were found in a gene, the most conserved one is shown. The network is grouped into 4 clusters based on protein-protein interaction score with connections across clusters shown by dashed lines. Genes not participating in a cluster were omitted for clarity. Genes in red were selected for rescue in this study. **(d):** *Left:* SNP reversal is carried out by targeting an sgRNA to a PAM (*NGG*, reverse strand displayed) proximal to the respective SNP (red). A ssDNA homology donor oligo carrying the reversed base (red) is provided as a repair template carrying additional, silent SNPs (green) to prevent re-targeting of the repaired sequence. *Right:* Sequence alignment of targeted SNP loci in *ATM* (R2830H, top) and *PRKDC* (D1641N, bottom). CHO-K1: host strain, Donor: homology oligo template, ATM+/PRKDC+: cell clones obtained from SNP reversal (PRKDC+ is short for ATM+ PRKDC+), C. gri: Chinese hamster (*Cricetulus griseus*). **(e):** Relative transcript abundance of *XRCC6* and *LIG4* in CHO-K1 WT (grey), CHO-K1 CMV::XRCC6 (light blue), and CHO-K1 CMV::LIG4 (dark blue). Welch’s t-test after log_2_-transformation, n=3.

### CHO cells contain numerous mutations in genome stability genes

To assess the extent of mutations in DNA repair genes, we analyzed whole-genome sequencing (WGS) data of 11 CHO cell lines, including those commonly used for cell line development in biopharmaceutical production (e.g. CHO-S, CHO-XB11, CHO-DG44) (Table 1). We extracted a list of genes related to genome stability from a curated database [50], and aligned the respective CHO homologs to the Chinese hamster genome [4, 5]. We looked for single-nucleotide polymorphisms (SNPs) and short indels (<10 bp), located in exons, and used a phylogeny-based score to predict the deleteriousness of these mutations on gene function (PROVEAN score) [51]. All 11 CHO lines were heavily affected by SNPs in DNA repair genes. We identified 157 mutations in various genes spanning 14 ontology classes (Fig 2a; Suppl Fig. 1). Using copy-number information derived from the WGS datasets, we retained only mutations which had undergone a loss of heterozygosity since these can typically be expected to have higher impacts on phenotype. After this filtering, the most conserved and most deleterious mutations occurred in genes related to non-homologous end-joining, DNA-damage sensing, and in helicases all of which are relevant for maintenance of genome stability (Fig. 2b).

**Table 1:** Genome sequences of CHO cell lines analyzed in this study.

| Cell line | Origin | NCBI Sequence Read Archive Number |
| --- | --- | --- |
| CHO-K1 | ATCC | SRP045758 |
| CHO-K1 | ECACC | SRS406579 |
| CHO-K1/SF | ECACC | SRS406580 |
| CHO protein-free | ECACC | SRS406578 |
| CHO-DG44 | Life Technologies | SRS406582 |
| CHO-S | Life Technologies | SRS406581 |
| CHO-S | Clone from the Technical University of Denmark<br>(derived from Life Technologies) | (unpublished) |
| C0101 | Undisclosed company (Drug producing cell line<br>derived from CHO-S from Life Technologies) | SRX258098 |
| CHO-Z | Clone from the Technical University of Denmark<br>(Serum-free suspension adapted clone derived<br>from an ECACC CHO-K1 clone) | (unpublished) |
| CHO-DXB11 | Clone from the Technical University of Denmark | SRX689758 |
| CHO-pgsA-745 | ATCC | (unpublished) |

### Restoration of DNA repair genes leads to an improved DNA damage response

Of the likely deleterious mutations (Fig. 2b), we selected critical DSB repair carrying conserved mutations across the cell lines in our dataset. In mammalian DNA repair, the protein kinase *Ataxia-telangiectasia mutated* (*ATM*) is among the first factors to become activated upon persistent DSBs that are not resolved by simple ligation, and instead require more advanced processing, such as end-resection, before repair can be attempted [52, 53]. *ATM* is thus a key upstream player in the DNA-damage response to DSBs (Fig. 2c) [54]. All 11 CHO cell lines carry a SNP in the catalytic PI3K/PI4K domain (R2830H), predicted to be highly deleterious according to PROVEAN scoring which made it a promising target for SNP correction. Downstream of *ATM*, DSB repair occurs through two main pathways: non-homologous end-joining (NHEJ) and homology-directed repair (HDR) [55]. In HDR, a sequence template homologous to the damaged strand is utilized to produce an error-free repair of the DNA lesion. In NHEJ, DNA ends from both sides of the lesion are ligated without using homology information which often implies loss of basepairs. While HDR is clearly preferable for maintaining sequence integrity, its activation requires homologous sequences which are only available during S- and G2-phase, but unavailable during the longer G1 phase. In addition, mutations in HDR genes were less widespread among CHO cell lines (Fig. 2b). Thus, we reasoned that restoration of NHEJ would be of primary importance to improve DNA repair in CHO and mitigate chromosomal instability and transgene loss. PRKDC is the catalytic subunit of the DNA-PK (DNA-dependent protein kinase) complex that becomes active early during NHEJ to shield loose chromosome ends at a break site and initiate ligation [56]. In our dataset, all 11 CHO cell lines carry two SNPs in *PRKDC*, both predicted to be detrimental. Thus, given its key role in NHEJ and this widespread occurrence, we reasoned that *PRKDC* would be another high-priority target for gene correction efforts. We prioritized SNP D1641N in the FAT domain over S3419G due its PROVEAN score indicating higher deleteriousness.

Using CRISPR/Cas9-mediated gene correction protocols (Integrated DNA Technologies), we generated a clonal CHO-K1 population with a successful reversal of R2830H in *ATM* (hereafter referred to as CHO-K1 ATM^+^). From this population, we generated a sub-clone with a successful reversal of D1641N in *PRKDC* (hereafter referred to as CHO-K1 ATM^+^ PRKDC^+^) (Fig. 2d). These reversals were done in succession to assess the cumulative effect of DNA repair improvements. RNA-seq of the new cell lines ATM^+^ and ATM^+^ PRKDC^+^ revealed only few differentially expressed genes, and gene set enrichment analysis did not identify significantly up-/downregulated pathways (Suppl. Fig. 2), consistent with these SNP reversals not having detrimental effects on viability or metabolism. In a parallel approach, we generated cell lines expressing intact copies of *XRCC6* and *LIG4*, cloned from Chinese hamster tissue (Fig. 2e. Hereafter referred to as CHO-K1 CMV::XRCC6 and CHO-K1 CMV::LIG4), which both carry likely detrimental SNPs (XRCC6 Q606H and LIG4 L145I and C741R). These genes were chosen despite the heterozygosity of these SNPs considering the critical roles these genes play in non-homologous end-joining [57].

To assess improvement in DSB repair capability, we implemented a GFP-based reporter system in CHO (based on the EJ5-GFP reporter developed for cancer cells [58]) that allows quantification of DSB repair through flow cytometry as sequence loss from compromised DSB repair causes GFP expression (Fig. 3a,b). We validated the assay in CHO-K1 wildtype cells treated with KU-60019, a highly effective small-molecule inhibitor against the ATM kinase, which caused a significant increase in the fraction of GFP+ positive cells, indicating compromised DSB repair (Fig. 3c). This also indicates that the R2830H mutation in *ATM* in CHO-K1 represents a hypomorphic allele since, otherwise, chemical inhibition of ATM would not have exacerbated the repair deficiency. Interestingly, CHO-K1 ATM^+^ showed a significant decrease in the fraction of GFP positive cells, indicating a successful improvement in repair of the induced lesion (Fig. 3c). Even further improvement of repair ability was seen in ATM+ PRKDC+ (Fig. 3c). CHO cells overexpressing intact *XRCC6* or *LIG4* copies also showed a reduction in GFP positive cells (Fig. 3d), indicating successful improvement in DSB repair ability in these cell lines.

**Fig 3:**
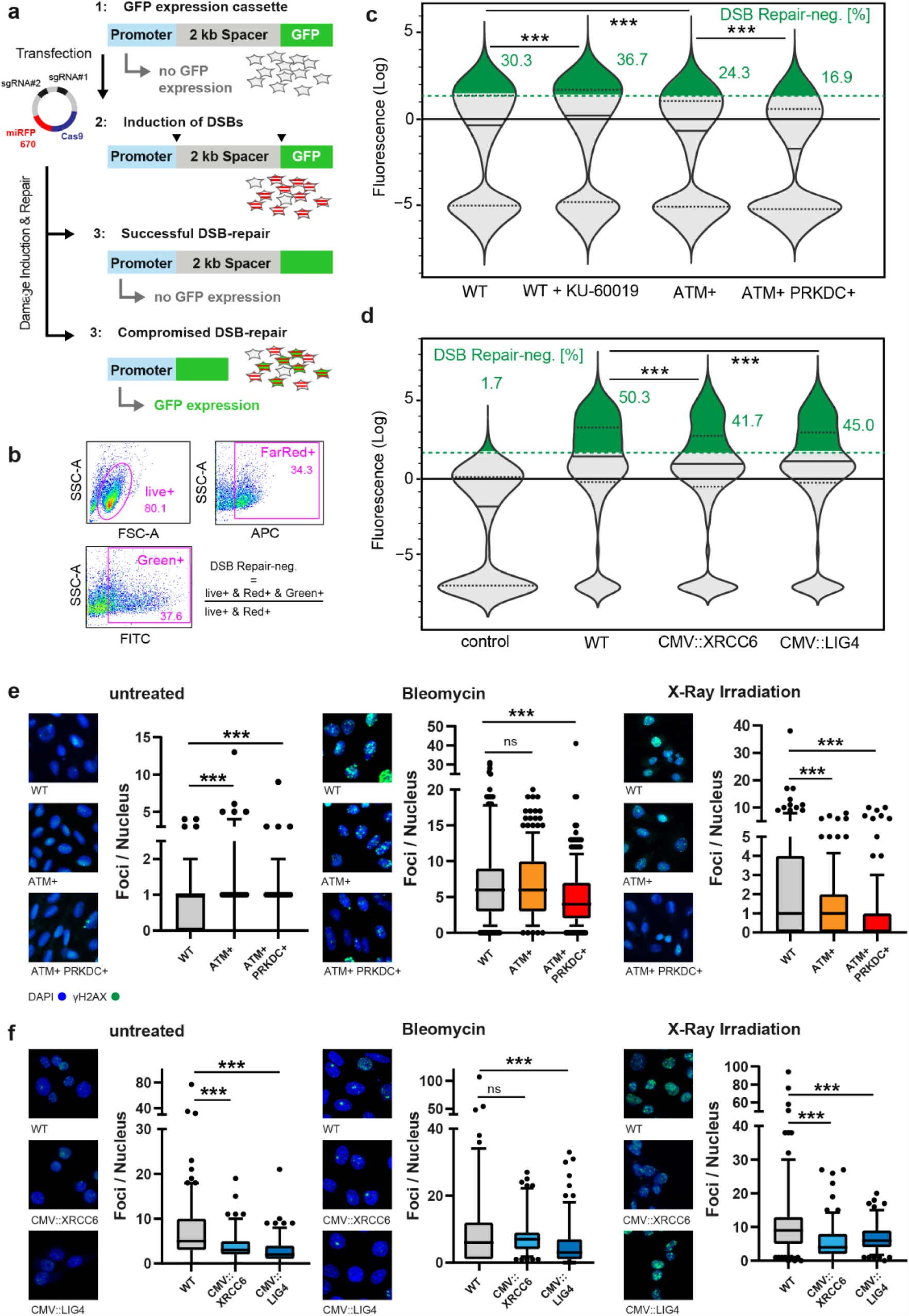
Analysis of DNA repair ability after gene rescue. **(a):** *Step 1*: The EJ5-GFP cassette comprises a promoter, a 2 kb spacer, and a GFP reading frame. The spacer prevents the promoter from driving GFP expression. *Step 2*: Transient transfection with a DSB-inducing plasmid, encoding Cas9 and two sgRNAs, targets two sites at the 5’ and 3’ ends of the spacer. Successfully transfected cells are identified through far-red fluorescence of the Cas9:miRFP670 protein. *Step 3*: Transfected cells that repair both DSBs properly keep the spacer in place and remain GFP-negative. Transfected cells that lose the spacer due to compromised DNA repair become GFP-positive (assay modified from [58]). **(b):** DSB repair ability is quantified through flow cytometry by relating the fraction of GFP-positive cells to all transfected cells, with the gates shown. **(c):** Violin plots of GFP signal in FarRed+ cells of the indicated genotypes. Data showing pooled populations from three independent transfections per genotype. Green dashed line: GFP intensity threshold. Two-sample Kolmogorov-Smirnov tests, n≥6,700 cells. **(d):** Violin plots of GFP signal in FarRed+ cells of the indicated genotypes. control: WT cells, not transfected with the DSB-inducing plasmid (not displayed in (c)). Data showing pooled populations from three independent transfections per genotype. Green dashed line: GFP intensity threshold. Two-sample Kolmogorov-Smirnov tests, n≥4,500 cells. Different percentages of GFP+ cells in WT cells across different experiments are caused by differences in transfection rates and recovery periods after transfection. **(e):** Immunostainings against γH2AX in CHO-K1 wildtype, ATM+, ATM+ PRKDC+. *Left*: untreated cells, n≥115 cells. *Middle*: Cells treated with 10 μg/mL bleomycin (24h recovery), n≥405 cells. *Right*: Cells treated with 0.5 Gy X-ray radiation (2h recovery), n≥196 cells. Welch’s t-tests. **(f):** Immunostainings against γH2AX in CHO-K1 wildtype, CMV::XRCC6, CMV::LIG4. *Left*: untreated cells, n≥140 cells. *Middle*: Cells treated with 10 μg/mL bleomycin (24h recovery), n≥114 cells. *Right*: Cells treated with 0.5 Gy X-ray radiation (2h recovery), n≥134 cells. Welch’s t-tests. Whiskers showing 5/95-quantiles. Cells counterstained with DAPI.

To rule out effects potentially specific to the described GFP reporter, we analyzed DSB repair efficiency more generally, through immunostaining against γH2AX. In CHO-K1, low levels of γH2AX foci are visible even in the absence of any DSB-generating treatments, corresponding to the endogenous origins of DSBs (Fig. 3e,f: left). We note that the generation of γH2AX is partially dependent on the ATM kinase [59] which likely explains why under untreated conditions foci numbers were slightly higher in cell lines carrying a restored *ATM* gene (Fig. 3e: left) as these cells likely mark damaged sites more effectively. In contrast, CMV::XRCC6 and CMV::LIG4 showed lower numbers of endogenous DNA damage foci (Fig. 3f: left). To compare repair efficiency after a strong DNA insult, we applied bleomycin treatment as described above, and found fewer foci in the ATM+ PRKDC+ and CMV::LIG4 cell lines, compared to CHO-K1 WT, after a 24h recovery period (Fig. 3e,f: middle). To rule out different rates of bleomycin uptake between cell lines [60], we also induced DNA damage with exposure to 0.5 Gy of X-ray irradiation. Foci enumeration after 2h of recovery showed improved DSB repair efficiency in all engineered cell lines compared to wildtype CHO-K1 (Fig. 3e,f: right). Thus, the DSB repair machinery is more active in the engineered cell lines and yields improved response to ubiquitous DNA damage, not specific to a break triggered at a specific site.

### Restoration of DNA repair improves genome stability in CHO

Unrepaired DSBs can lead to chromosomal aberrations, ultimately driving loss of transgene expression. We thus asked whether the improvements in the DNA damage response in the engineered CHO cell lines would improve the overall state of genome integrity. For this, we first exposed wildtype and engineered cell lines to DSB-inducing conditions and analyzed genome integrity on the single-cell level by electrophoresis where both the length and the intensity of the resulting DNA tail is an indicator of the amount of genome fragmentation (comet assay). After exposing cells to bleomycin or radiation, we noticed considerable DNA displacement in wildtype CHO cells, with some cells exhibiting very long, bulky DNA tails indicating severe genome fragmentation due to persistent DSBs (Fig. 4a,b: middle and right). Restoration of *ATM* resulted in minor changes in DNA tail length, but additional restoration of *PRKDC* led to a strong reduction in both tail length and intensity, and we did not detect long bulky DNA tails in these samples (Fig. 4a: middle and right). These results are consistent with *ATM* acting as a DNA damage sensor [54], not a bona fide DNA repair gene, which likely explains why rescue of *ATM* alone had limited effects. Importantly, also in the absence of genotoxic stress, ATM*+* PRKDC*+* showed very short DNA tails, indicating improved genome stability (Fig. 4a: left). We also observed shorter DNA tails in CMV::XRCC6 and CMV::LIG4 cell lines across all conditions (Fig. 4b). Together, these results indicate that restoration of key genes controlling non-homologous end-joining successfully enhances DNA repair and reduces genome fragmentation. We noticed several chromosome aberrations in the parental CHO-K1 strain, from which the engineered cell lines were derived, such as translocations, e.g. on chromosomes #3, #6, or #7, and whole chromosome duplications, e.g. #4 and loss of X-chromosomes (Fig. 4c). In a long-term culture experiment, where cell lines were passaged in parallel over a 60-day period, ATM+ and ATM+ PRKDC+ cell lines acquired considerably fewer additional chromosome aberrations than the parental wildtype cell line (Fig. 4d,e), consistent with improved genome stability. Another wildtype sample, supplemented with 3 μM KU-60019, served as a positive control and acquired many more aberrations, resulting in almost 75% of karyotypes deviating from the parental main karyotype (Fig. 4d,e). In summary, while CHO cells carry a high burden in DNA repair genes, restoration of just few key genes can improve DSBs repair as well as structural chromosomal stability.

**Fig 4:**
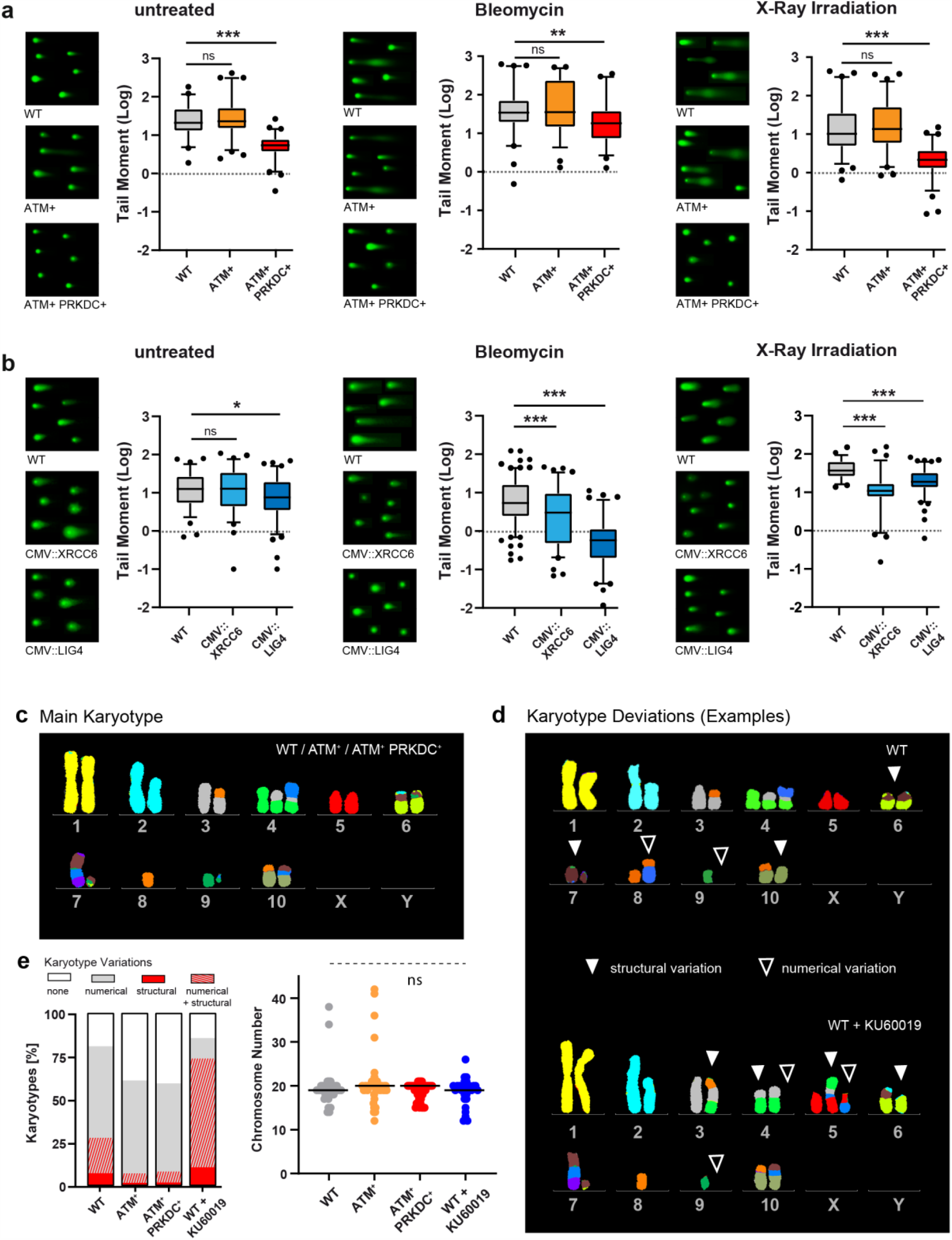
Analysis of genome stability after gene rescue. **(a):** Representative composite images and tail moments (=tail length*DNA in tail [%]) of wildtype, ATM+ and ATM+ PRKDC+ cells after electrophoresis in a low-melting agar (comet assay). *Left*: untreated cells, n≥53 cells. *Middle*: Cells treated with 10 μg/mL bleomycin (24h recovery), n≥56 cells. *Right*: Cells treated with 0.5 Gy X-ray radiation (2h recovery), n≥71 cells. Welch’s t-tests. Whiskers showing 5/95-quantiles. **(b):** Representative composite images and tail moments (=tail length*DNA in tail [%]) of wildtype, CMV::XRCC6 and CMV::LIG4 cells. *Left*: untreated cells, n≥74 cells. *Middle*: Cells treated with 10 μg/mL bleomycin (24h recovery), n≥86 cells. *Right*: Cells treated with 0.5 Gy X-ray radiation (2h recovery), n≥57 cells. Welch’s t-tests. Whiskers showing 5/95-quantiles. **(c):** Main karyotype after 60 passages of long-term culture. Chromosomes were identified using pseudo-color probes, specific for each *Cricetulus griseus* chromosome. **(d):** Examples for deviating karyotypes in WT (top) and WT, supplemented with KU-60019 (bottom). Open arrows indicate a numerical variation (i.e. gain/loss of a chromosome), closed arrows indicate a structural variation (i.e. an altered color pattern). **(e):** *Left*: Classification of karyotypes into: (i) showing at least one numerical variation with no structural variations (grey), (ii) showing at least one structural variation with no numerical variations (red), (iii) showing both at least one numerical and at least one structural variation (grey/red striped), and (iv) showing no variations (white), relative to the main karyotype (c). Differences in frequency of structural variations (red and red/grey fractions) significant at 5% level (Binomial test) (asterisks omitted for clarity). Averaged fractions from duplicate experiments: WT n=26/34; ATM+ n=21/37; ATM+PRKDC+ n=21/37; WT+KU60019 n=8/19. *Right:* Total number of chromosomes per karyotype. Bar = median. Non-parametric ANOVA (Kruskal-Wallis test).

### Restoration of DNA repair improves titer stability in a producing cell line

Based on the notion that genome instability can disrupt the maintenance of high protein titers in industrial biomanufacturing, we reasoned that genome stabilization could counteract this problem by slowing the loss of transgene copies caused by chromosome instability. The results above support the notion that restoration of DNA repair genes could help stabilize protein titers. Thus, we applied this strategy in CHO-SEAP, an adherent cell line expressing human secreted alkaline phosphatase (SEAP) [61]. Guided by our results for non-producing CHO-K1, we generated CHO-SEAP cell lines expressing Chinese hamster wildtype copies of *XRCC6* and *LIG4*, respectively. These cell lines, CHO-SEAP CMV::XRCC6 and CMV::LIG4, showed significantly improved DSB repair ability, evident as a reduction of GFP positive cells in the EJ5-GFP assay (Fig. 5a) and in DNA repair foci (Fig. 5b). Surprisingly, reversals of *ATM* R2830H and *PRKDC* D1641N in CHO-SEAP CMV::XRCC6 did not yield further improvements, but instead caused a decrease in DSB-repair ability (Fig. 5a). Consistent with this observation, chemical inhibition of ATM resulted in a slight improvement in repair ability (Fig. 5a), in contrast to our observations in CHO-K1 (See Discussion).

**Fig 5:**
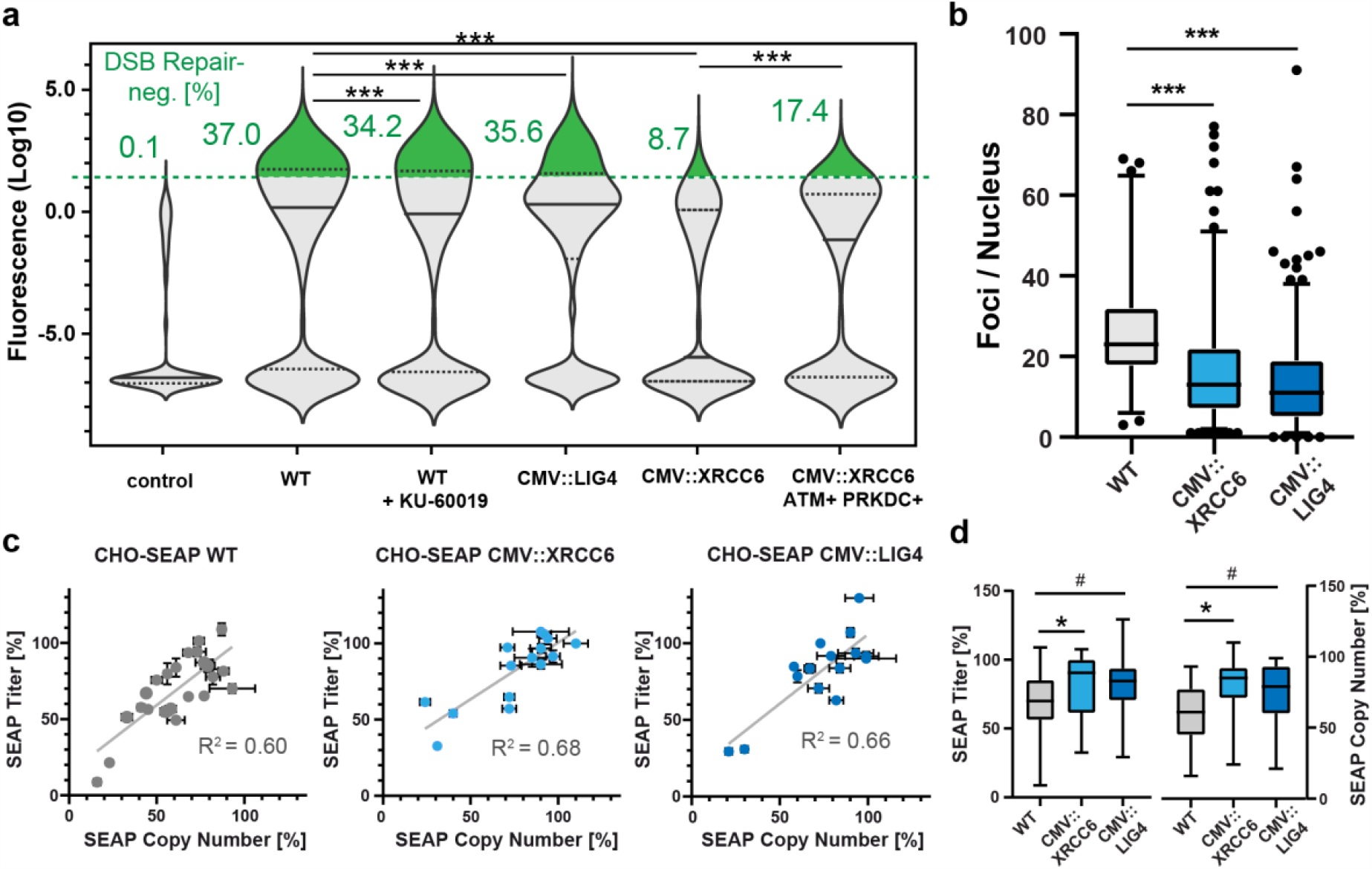
DSB repair and protein titer in a producing CHO cell line. **(a):** Violin plots of GFP signal in FarRed+ cells of the indicated genotypes in CHO-SEAP. control: WT cells not transfected with the DSB-inducing plasmid. Data showing pooled populations from two independent transfections per cell line. Green dashed line: GFP intensity threshold. Two-sample Kolmogorov-Smirnov tests. n≥3,800 cells. **(b):** Immunostainings against γH2AX in CHO-SEAP wildtype, CMV::XRCC6, and CMV::LIG4. Cells treated with 10 μg/mL bleomycin (24h recovery). Pooled data from 4 representative clones per genotype. n≥134 cells. Welch’s t-tests. Whiskers showing 2.5/97.5-quantiles. **(c):** SEAP titer change versus SEAP gene copy number change in clonal populations of the indicated genotypes over a long-time culture period of 74 days. Percentages indicate retention of SEAP titer and copy number at day 74 relative to day 0. **(d):** Boxplots of the data shown in (c). Welch’s t tests. WT: n=23, CMV::XRCC6: n=15, CMV::LIG4: n=16. Whiskers showing min/max.

Because both *XRCC6* and *LIG4* overexpression had beneficial effects on genome stability in CHO-SEAP, we next grew clonal populations of CHO-SEAP WT, CMV::XRCC6, and CMV::LIG4 in parallel over a 74-day period and compared both SEAP titer and SEAP transgene copy number at the beginning and the end of the experiment. Prior to the start of the experiment, cells were cultured in 5 μM methotrexate (MTX) for 1 week to select for high SEAP copy number amplification, after which MTX was taken off the growth medium. MTX is a competitive inhibitor of dihydrofolate reductase, an essential metabolic enzyme, which is co-expressed with the transgenic SEAP locus [61]. In all cell lines, SEAP titer retention (=titer at day 74, relative to day 0) correlated significantly with SEAP copy number retention (Fig. 5c), highlighting the importance of genome stability as a determinant of cell line productivity. CHO-SEAP WT cells retained about 75% of the initial SEAP titer on average, but a considerable fraction of clones showed dramatic titer decline to as low as 10% (Fig. 5c: left). Clones overexpressing *LIG4* showed improvements in stability of both titer and copy number although results were only marginally statistically significant (Fig. 5c: right; Fig. 5d). *XRCC6* overexpression, however, gained significant stabilization of both SEAP titer and copy number, retaining about 90% SEAP titer on average, and showing fewer cases of dramatic titer decline, with the minimum titer retention at 30% (Fig. 5c: middle; Fig. 5d). Together, these results show that rescue of non-homologous end-joining in CHO cells successfully mitigates transgene loss and titer decline.

## Discussion

The CHO-based platform in biologics manufacturing is well-established, but an important unsolved problem is genome instability, which causes a population derived from a clonal cell to display genetic heterogeneity. Apart from few studies identifying impaired repair pathways [62, 63], this is the first report documenting the full extent of mutations affecting DNA repair genes across CHO cell lines. Moreover, while reactivation of silenced DNA repair genes has been successfully implemented before [64], restoration of DNA repair ability has not yet been systematically explored to mitigate genome instability in cell line development. This study is the first report to show that restoring DNA repair function through genome editing ameliorates genome stability in CHO. Furthermore, we showed that despite the high mutational burden in DNA repair genes, restoration of just a single gene can yield significant improvements in genome integrity. This makes DNA repair restoration a powerful and feasible novel addition to the cell line engineering toolbox. Our dataset of mutated DNA repair genes opens up a plethora of options for future studies, targeting single genes or combinations of genes to develop novel cell lines for biopharmaceutical manufacturing. While effective alternative approaches have recently been described to increase CHO cell productivity, such as overexpression of key metabolic genes [65], suppression of apoptosis [13], or design of novel promoters [66], restoration of DNA repair tackles the mechanistic root of genome instability and could enable long-lasting stability improvements. Beyond protein expression, restoration of DNA repair genes could prove effective in other aspects of cell line engineering, for example by improving rates of targeted gene integration or gene correction in CHO [67]. Also, the approach could be expanded to other cell lines.

As shown here, improvement of DSB repair ability appears to occur in an incremental fashion when combinations of DNA repair genes are being restored, provided these genes work synergistically, as ATM and PRKDC do in CHO-K1, for example, evident from the gradual improvement in repair ability when combining both gene corrections (Fig. 3c). However, selecting the optimal gene targets, or combination of gene targets, for DNA repair rescue remains challenging. While data on human cancers, DNA repair, or evolutionary conservation [68] can guide selection of likely effective candidate genes, our data shows that restoration of even the same genes can yield different outcomes in different cell lines (Fig. 5a). Given the divergent genomes of different CHO cell lineages [5] and the complex, intertwined nature of the mammalian DSB repair cascade [57], results from one cell line may not necessarily apply likewise to others. In mammals, DSB repair follows a “decision tree” [57] where pathway choice is largely determined by the severity of the DNA lesion. In particular, while a core NHEJ pathway can act independently of ATM [57, 69], ATM plays a key role in initiating repair of lesions requiring more pre-processing and more advanced repair pathways, such as homology-directed repair (HDR), alternative end-joining (aEJ), or the Fanconi anemia (FA) pathway [57, 70, 71]. For this repair system to be effective, genes in these pathways downstream of ATM need to be functional. Thus, it is possible that in CHO-K1 these pathways have retained higher functionality than in CHO-SEAP which would explain why activating ATM in CHO-SEAP has detrimental outcomes. Indeed, our dataset shows a higher incidence of SNPs in HDR or FA pathways in CHO-SEAP (a DXB11 derivative) compared to CHO-K1 (Suppl. Fig. 1). Thus, in CHO-SEAP, *ATM* restoration might have triggered a negative net effect with downstream pathways being largely incapacitated, especially since the competition between pathways could inhibit functional NHEJ [72]. Previous studies reported similar unexpected effects upon inhibition of DNA repair genes, such as *ATM* or *MRE11* [67, 73]. Observing such opposite effects in different CHO cells after rescuing identical genes provides a promising model platform to study synergistic gene relationships and competition within the DSB repair hierarchy. Unlike *ATM*, restoration of *XRCC6* resulted in a considerable improvement in DSB repair in both CHO-K1 and CHO-SEAP, although the SNP (Q606H) in *XRCC6* is only heterozygous. Yet, Ku70 (the protein encoded by *XRCC6*) binds to Ku80 to form the heterodimeric Ku complex, and mutations in *XRCC6* are thus likely to cause a dominant phenotype. Indeed, in human cells, a heterozygous Ku80 mutation can trigger increased genome instability [74]. Thus, target choice for gene correction needs to be carefully considered. Developing a high-throughput screening method will therefore be highly valuable to identify optimal gene targets. The EJ5-GFP cell line described in this study can serve as an excellent discovery tool that could be utilized in flow-cytometry screens to identify optimal targets for DNA repair enhancement for this purpose. Certainly, the EJ5-GFP assay is approximate due to the possibility of false negative signal (i.e. a reporter site that did not get cut despite the presence of Cas9:miRFP, or a reporter site whose loose ends failed to merge entirely), but it still provides a good estimate of DSB repair ability since positive GFP expression can only occur after imperfect DSB repair processing. Further, cell line stabilization efforts targeting enhancement of DNA repair have to occur late in the cell line development process since transgene copy number amplification requires high genome plasticity. Thus, for optimal efficacy, genome stabilization preferably needs to occur after a high-producing clone has been isolated.

In conclusion, this study provides the first thorough documentation of the deficient DNA repair machinery in CHO cells, and constitutes a proof-of-concept of the notion of DNA repair restoration as a powerful novel method for cell line development in industrial protein expression to ensure stable upstream manufacturing processes.

## Supplemental Figures

**Suppl. Fig 1:**
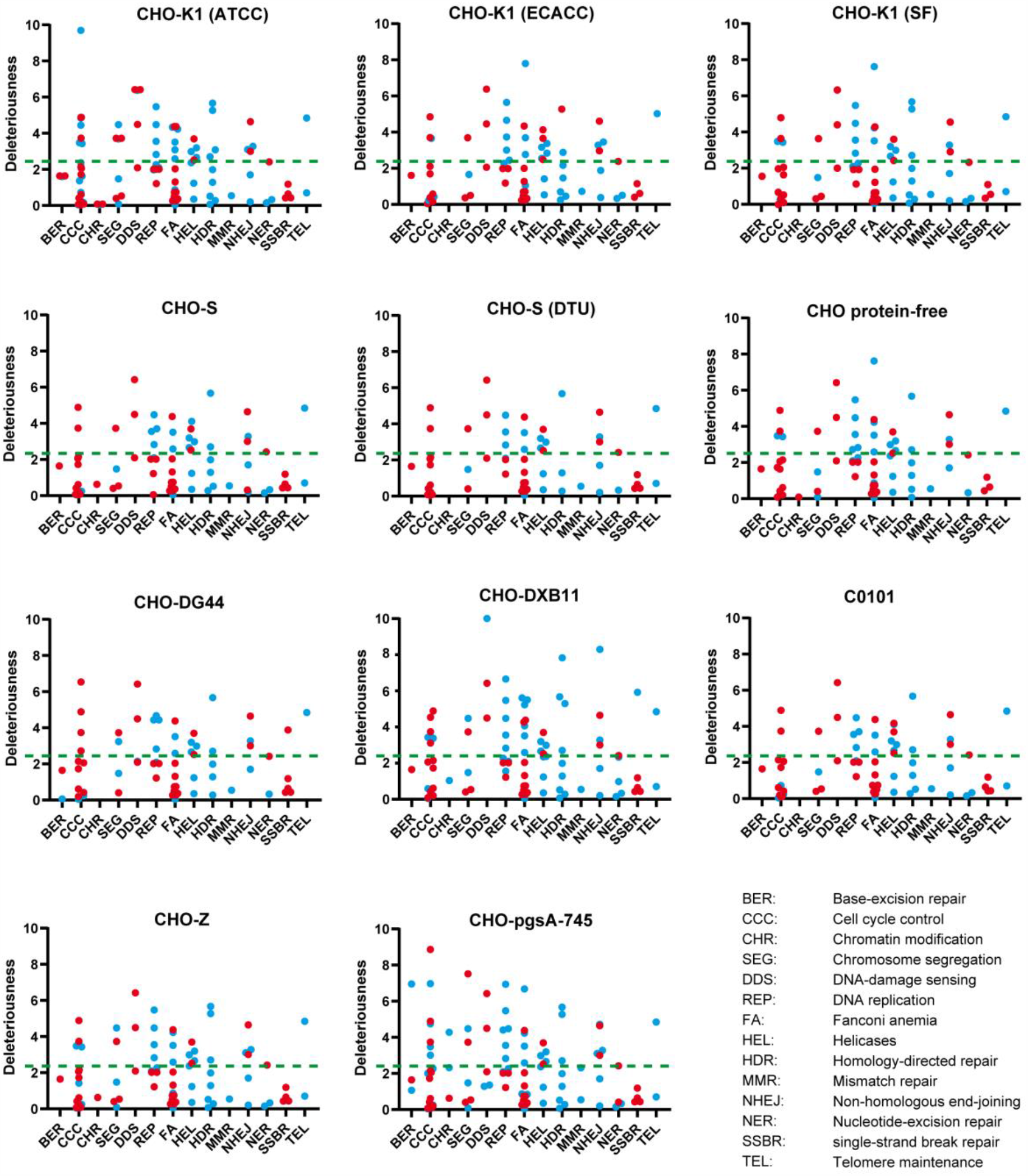
Mutations in genes relevant to genome stability in various CHO cell lines. Predicted deleteriousness (Negative PROVEAN score) of mutations (SNPs and Indels) in genes related to genome stability (see legend at bottom-right) in a panel of commonly used CHO cell lines. Blue dots indicate heterozygous mutations, red dots indicate loss of heterozygosity. Green dotted line indicates significance threshold [51].

**Suppl. Fig 2:**
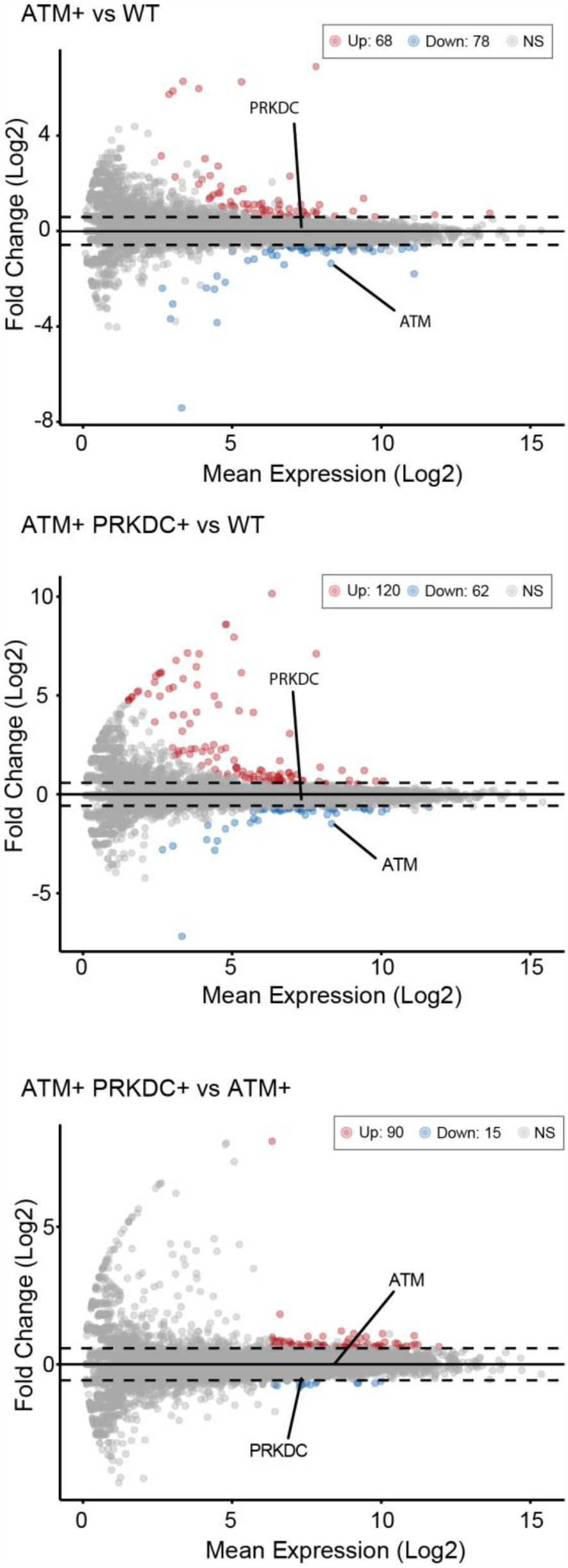
Transcriptomic analysis of CHO-K1 cell lines after SNP reversals. *Top*: Transcript abundance (x-axis) in CHO-K1 ATM+. Transcript fold-change (y-axis) relative to CHO-K1 WT. *Middle*: Transcript abundance (x-axis) in CHO-K1 ATM+ PRKDC+. Transcript fold-change (y-axis) relative to CHO-K1 WT. *Bottom*: Transcript abundance (x-axis) in CHO-K1 ATM+ PRKDC+. Transcript fold-change (y-axis) relative to CHO-K1 ATM+. Significantly up-/downregulated genes highlighted in red/blue.

## Materials & Methods

### Sequence analysis and statistics

Whole-genome sequencing data of 11 CHO cell lines [4, 5] (Table 1) was downloaded from Genbank (https://www.ncbi.nlm.nih.gov/sra), together with the Chinese hamster genome [5]. Raw sequencing reads were pre-processed using FastQC [75] for quality control and Trimmomatic [76] to remove low-quality base pairs and adapters. For mapping, the RefSeq CHO genome assembly GCF_000223151.1 was used. Transcript sequences of DNA repair genes [50] extracted from the reference genome assembly are known to have a low proportion of mismatches compared to the higher quality RefSeq transcripts. Thus, to ensure the reference genome has the correct sequences of DNA repair genes, RefSeq transcript sequences of DNA repair genes were merged into one scaffold separated by a 500 letter N gap. In cases of multiple isoforms, the transcript with the longest coding sequence was selected. Genomic coding regions of corresponding DNA repair genes were masked. Then the sequencing reads were mapped against the new reference genome using BWA [77]. Non-synonymous SNPs and InDels (including heterozygosity) were identified using the *gatk3*.5 software package [78] using standard parameters and annotated using SnpEff [79]. SnpSift [80] was used to filter genes with ontologies related to DNA repair [50]. Deleteriousness of mutations was predicted based on the PROVEAN score [51]. Protein-protein interaction networks were generated with STRING [81]. RNA was extracted in from triplicate flasks (RNeasy, Qiagen) and stored at −80°C before sequencing. RNA-sequencing data quality was assessed using FastQC [75]. Adapter sequences and low-quality bases were trimmed using Trimmomatic [76]. Reads were then quasi-mapped to the CHO-K1 genome [4, 5] and quantified with Salmon [82] using default parameters. Raw gene expression data were then normalized and analyzed for differential expression using DESeq2 [83]. After Benjamini-Hochberg FDR correction, genes with adjusted p-values less than 0.05 and fold changes greater than log_2_(1.5) were considered differentially expressed. Statistical tests and sample sizes are indicated in the figure legends. *** p<0.001, ** p<0.01, * p<0.05, # p<0.1, ns not significant. Tests were carried out in Prism (GraphPad) or R (The R Project for Statistical Computing).

### Cell culture and cell line generation

CHO-K1 cells (ATCC: CCL-61) and CHO-SEAP cells [61] were cultured in F-12K medium (Gibco), or Iscove’s Modified Dulbecco’s Medium (IMDM), respectively, supplemented with 10% (v/v) fetal bovine serum (FBS, Corning) and 1% (v/v) penicillin/streptomycin (Gibco) at 37°C under an atmosphere of 5% CO_2_. Cells were passaged every 2-3 days. CHO-K1 EJ5-GFP and CHO-SEAP EJ5-GFP were generated by transfecting CHO-K1 cells, or CHO-SEAP cell respectively, with a XhoI-linearized EJ5-GFP plasmid (Addgene #44026) and subsequent combined selection with puromycin (7 μg/mL) and hygromycin (300 μg/mL). After two weeks of antibiotic selection, clonal populations were generated by seeding cells in limiting dilution on 96-well plates and visually selecting clonal colonies. EJ5-GFP insertion was verified by PCR (OneTaq, New England Biolabs). CHO-K1 ATM+ was generated by transfecting a clonal population of CHO-K1 EJ5-GFP with a Cas9:tracrRNA:sgRNA ribonucleoprotein particle (Integrated DNA Technologies), targeting R2830H in ATM (Gene ID: 100754226), and a homology donor oligo encoding the corrected sequence, following standard protocols (Integrated DNA Technologies). Clonal populations were generated through limiting dilution, and the R2830H site was screened by PCR for the presence of a TaqI site in the corrected locus and verified by Sanger sequencing (Eton Biosciences). Sanger sequencing data was deconvoluted using the ICE Analysis Tool (Synthego). CHO-K1 ATM+ PRKDC+ was generated by transfecting a clonal population of CHO-K1 ATM+ with a Cas9:tracrRMA:sgRNA ribonucleoprotein particle, targeting D1641N in PRKDC (Gene ID: 100770748), and a homology donor oligo encoding the corrected sequence. Clonal populations were generated through limiting dilution, and the PRKDC D1641N site was screened by PCR for the presence of a BamHI site in the corrected locus and verified by Sanger sequencing. CHO-SEAP CMV::XRCC6 and CHO-SEAP CMV::LIG4 were generated by transfection of CHO-SEAP EJ5-GFP with the corresponding expression plasmids and maintained in IMDM media. At 24 hours post transfection, single cell clones were generated by serial dilution in 96-well plates and maintained in growth media with 200 μg/mL Zeocin and 5 μM methotrexate (MTX, EMD Millipore). CHO-SEAP CMV::XRCC6 ATM+ PRKDC+ was generated analogously to CHO-K1 ATM+ PRKDC+. Transfections with ribonucleoprotein particles were carried out using a Neon electroporation system (ThermoFisher) (12-well format). Transfection of DSB repair gene plasmids was carried out using a Nucleofector Kit T (Lonza) with 6×10^6^ cells transfected and 6 μg plasmid. All cells were maintained under combined puromycin/hygromycin selection throughout the experiments to avoid loss of the EJ5-GFP insertion. Double-strand-breaks were induced using bleomycin sulfate (Sigma-Aldrich). ATM was inhibited with KU-60019 (Selleckchem).

### Cloning of Chinese hamster genes

Chinese hamster (*Cricetulus griseus*) tissue (lung, liver) was a gift from George Yerganian. RNA extraction (RNeasy, Qiagen) and total cDNA synthesis (SuperScriptIII, Invitrogen) were carried out using standard protocols. cDNA was purified and concentrated using ethanol precipitation, and 1 μL purified total cDNA (100-200 ng) was used to amplify target genes through high-fidelity PCR (Q5, New England Biolabs). A vector backbone fragment was obtained by PCR amplification from plasmid pcDNA3.1/zeo(+) (ThermoFisher). Plasmids expressing DSB repair genes were constructed via Gibson assembly of gene fragment(s) and the vector fragment following the manufacturer’s instructions (New England Biolabs).

### EJ5-GFP flow cytometry assays

The DSB-inducer plasmid was constructed by ligation of two sgRNAs, targeting the EJ5-GFP cassette, into pX333 (Addgene #64073), and subsequent DrdI/KpnI-subcloning of the entire dual sgRNA expression cassette into pSpCas9(BB)-2A-miRFP670 (Addgene #91854). To run the EJ5-GFP assay, cells were transfected with 1 μg of this plasmid (Lipofectamine LTX, Invitrogen; 12-well format). After 30h cells were trypsinized, resuspended in 250 μL DPBS (Gibco), and analyzed on a Canto II flow cytometer (BD Biosciences). Untransfected cells served as negative control to define proper gates in the APC and FITC channels for miRFP and GFP expression, respectively. DSB-repair negative cells were identified through Boolean gating, as shown in Fig. 3b. Flow cytometry data was analyzed in FlowJo (BD Biosciences) and Prism (GraphPad).

### Immunofluorescence, comet assays and microscopy

Cells were seeded on chambered slides (Nunc, ThermoFisher) and, after attachment, either treated with 0.5 Gy X-ray radiation (X-RSD 320, Precision X-ray), or incubated with 10 μg/mL bleomycin sulfate (Sigma-Aldrich) for 12h. After the indicated recovery time, cells were fixated in 4% paraformaldehyde (ThermoFisher) for 10 min, washed in PBS (Gibco) for 2 min, and permeabilized with 0.5% Triton-X (Amresco) for 5 min, followed by washing for 5 min in PBS. After blocking with 5% goat-serum (MilliporeSigma) for 1h, cells were incubated in anti-γH2AX antibody (Cell Signaling Technology, Rabbit #9718) at 1:1000 dilution for 1h, washed three times in PBS-T (=0.1% Triton-X in PBS) for 5 min, and incubated with DyLight 488 goat-anti-rabbit (ThermoFisher) for 1h in the dark. After three washes in PBS-T for 5 min, cells were mounted in anti-fade mounting medium, containing DAPI (Vectashield Vibrance, Vector Laboratories). Samples were analyzed on a SP8 confocal microscope (Leica) with identical settings for gain and offset for each sample. Images were analyzed using ImageJ (NIH), following standard protocols for foci enumeration (https://microscopy.duke.edu/guides/count-nuclear-foci-ImageJ). Comet assays were carried out following the manufacturer’s protocol (Abcam), with 45 min electrophoresis at 1 V/cm in TBE-buffer. Nuclei were stained with Vista DNA Green (Abcam). Slides were analyzed on an Axio Imager 2 (Zeiss) and processed using the OpenComet plug-in (www.cometbio.org/index.html) for ImageJ (NIH).

### Long-term culture and SEAP quantification

For long-term culture, selected cell lines were seeded at 1×10^5^ cells/mL and subcultured for four passages with 5 μM MTX in 24-well plates. Then, cells were subcultured every three or four days for 74 days without MTX selection. Cell lines expressing *XRCC6* and *LIG4* were continuously maintained in media with 200 μg/mL Zeocin. Cell lines at Day 0 and Day 74 were seeded at 1×10^5^ cells/mL in 6-well plates containing 3 mL growth media per well without MTX. When the cell density reached ∼80% confluency, culture supernatant samples were collected for SEAP quantification (PhosphaLight, AppliedBiosystems). Alkaline phosphatase from human placenta (Sigma-Aldrich) was assayed and used as a standard. Gene copy number was determined by Real-Time PCR using TaqMan Fast Advanced Master Mix (Thermo Fisher) performed on a 7500 fast system (Applied Biosystems) according to the manufacturer’s protocol. PCR primers and TaqMan probes were designed and synthesized by IDT (Integrated DNA Technologies). Genomic DNA was extracted from Day 0 and Day 74 cell samples (DNeasy, Qiagen). The diluted genomic DNA of one CHO-SEAP clone (400 ng, 200 ng, 100 ng, 50 ng, 25 ng, 0 ng) was used to make a standard curve. Relative SEAP gene copy number in each cell sample was quantified in duplicate using 100 ng genomic DNA as a template with β-Actin as a reference.

### Karyotype analysis

Metaphase spreads were prepared as previously described [37]. Samples were labeled with multi-color DNA fluorescence in situ hybridization (FISH) probes (12XCHamster mFISH probe kit, MetaSystems) for spectral karyotyping as previously described [84]. For karyotypic analyses, the most abundant karyotype across samples was defined as the representative (“main”) karyotype, and deviations from this karyotype were scored as a numerical alteration (whole-chromosomal aneuploidy) and/or structural alteration (inter-chromosomal rearrangement, visible deletion). Structurally aberrant karyotypes (Fig. 4d) were defined as karyotypes showing at least one structural deviation from the representative karyotype.

### DNA oligos

#### Primers

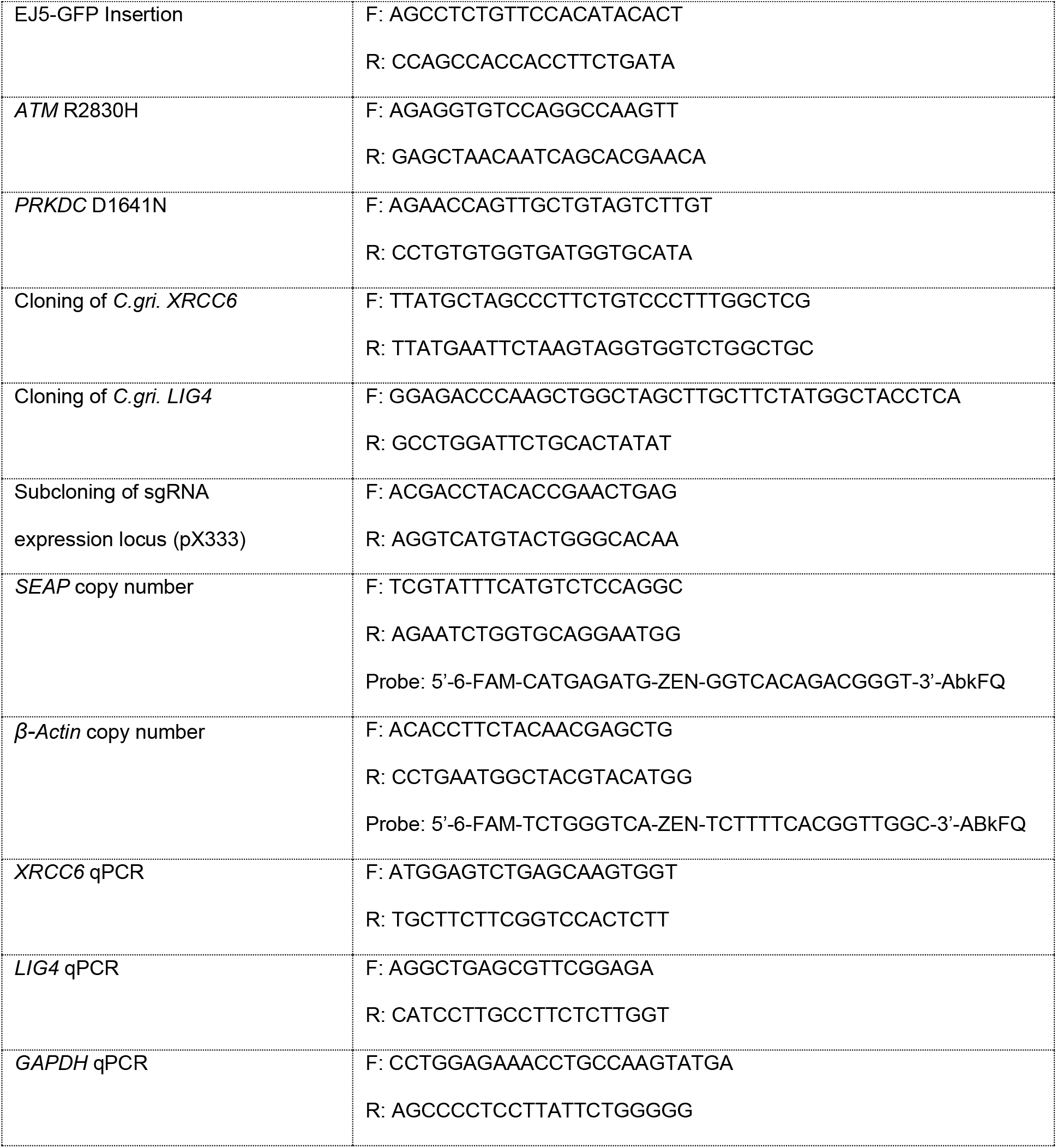

All primers were designed using Primer3 [85].

#### sgRNAs

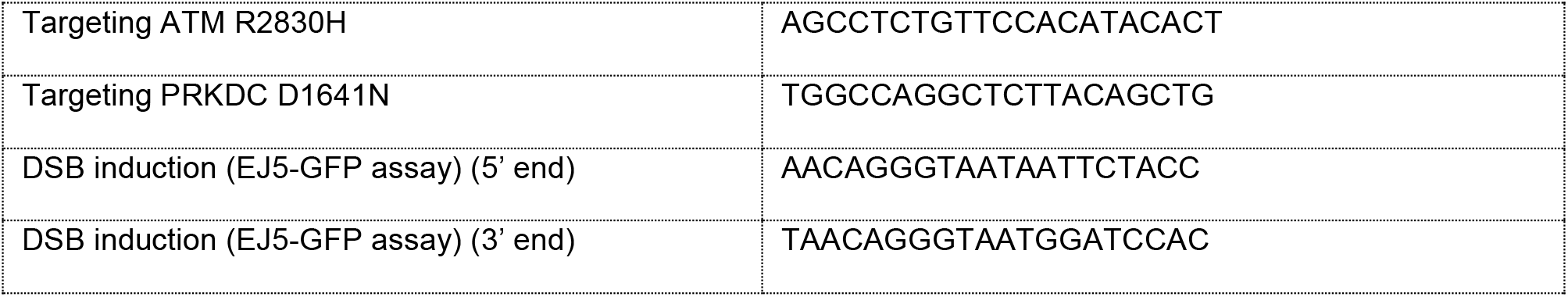

#### ssDNA Oligos

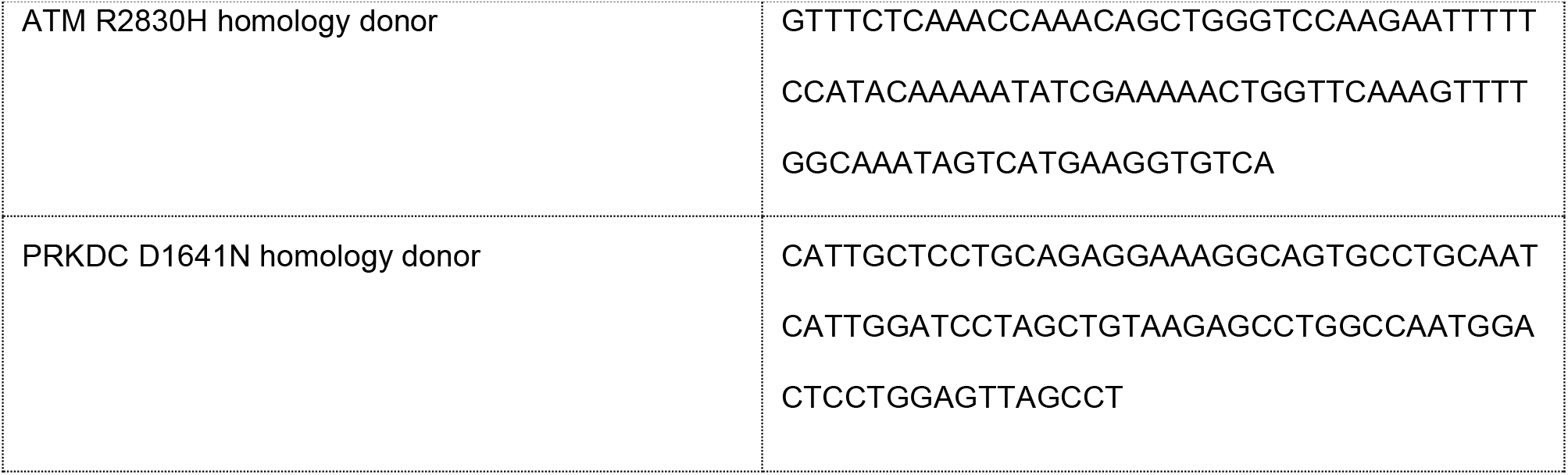

## Acknowledgements

We would like to express our deep gratitude to George Yerganian for sharing primary Chinese hamster tissue. We thank Bjørn Voldborg and Alexandra Hoffmeyer for sharing cell lines and for RNA sequencing services. We would also like to thank Francisco Diaz, Jennifer Santini and Marcy Erb for technical assistance, and Nicole Borth for sharing protocols. We also thank John Ruano-Salguero for revising the manuscript, Jong Youn Baik, and Philip Gordts for constructive feedback, and John Koh and April Kloxin for sharing cell lines. This work was supported by NIGMS (R35 GM119850, NEL), the Novo Nordisk Foundation (NNF10CC1016517, NEL) and NSF (grants 1736123 and 1412365, KHL). Laser-scanning confocal microscopy was supported by NINDS P30 NS047101.

## Author Contributions

PNS, XZ, QH, NKH, PL, KHL, NEL designed research; PNS, XZ, QH, HH, SL, CCK, YH, JCL performed research; PNS, XZ, QH, HH, SL, CCK, YH, JCL, PL, KHL, NEL analyzed data; PNS, XZ, KHL, NEL wrote the paper.

